# Causes of death in newborn C57BL/6J mice

**DOI:** 10.1101/2020.02.25.964551

**Authors:** Sara Capas-Peneda, Gabriela Munhoz Morello, Sofia Lamas, I Anna S Olsson, Colin Gilbert

**Affiliations:** Laboratory Animal Science, IBMC - Instituto de Biologia Molecular e Celular, Universidade do Porto, Rua Alfredo Allen 208, 4200-135 Porto, Portugal (SCP, GMM, IASO); i3S - Instituto de Investigação e Inovação em Saúde, Universidade do Porto, Rua Alfredo Allen 208, 4200-135 Porto, Portugal (SCP, GMM, SL, IASO); Animal Facility, IBMC - Instituto de Biologia Molecular e Celular, Universidade do Porto, Rua Alfredo Allen 208, 4200-135 Porto, Portugal (SL); Babraham Institute, Babraham Hall, Cambridge, CB22 3AT, United Kingdom (CG)

**Keywords:** Neonatal mortality, Mouse, causes of death, C57BL/6J

## Abstract

Neonatal mortality in wild-type laboratory mice is an overlooked welfare and financial problem in animal facilities around the world. Causes of death are often not reported and its causes remain unknown.

In this study, 324 newborn pups from two breeding colonies of healthy wildtype C57BL/6 mice underwent post-mortem analysis with special focus on obtaining proof of life after birth, evaluation of stomach contents and observation of congenital abnormalities that could compromise survival.

Based on a combination of lung morphology findings, outcome of lung float test, stomach contents and brown adipose tissue colouration, 21.6% of the pups found dead were considered stillbirths. Of the livebirths, only 3.2% were observed to have milk inside the stomach, indicating successful suckling. Congenital abnormalities were diagnosed only in a small fraction of the pups analysed. These results suggest that starvation was the most common cause of death, followed by stillbirth.

---

High rates of pre-weaning mortality in laboratory mice are a widespread issue among animal facilities worldwide. This represents a major welfare problem^33^, a significant financial burden through lowered productivity and has an impact on the 3Rs by increasing the number of animals needed and used for breeding.

Rates of perinatal mortality are reported as higher in inbred strains, especially in C57BL/6 mice, where they can reach 50%^19^. Attention to morbidity and mortality in newborn mice occurs primarily in the context of phenotyping genetically modified mice^2, 29, 45^. In contrast, knowledge about causes of neonatal death in breeding colonies of healthy wildtype mice is limited and currently there are no agreed published guidelines or strategies to address this issue.

Mice are an altricial species and pups are dependent on parental care within a maternal nest during the first phase of their lives. The pups are born helpless, relatively immobile, blind, hairless, capable of suckling but not able to otherwise fend for themselves. They possess a very limited ability to produce heat by non-shivering thermogenesis using brown adipose tissue (BAT)^6, 18^. Consequently, major causes of neonatal mortality and low viability could be related to failures of parental care, such as predation, starvation and hypothermia in addition to birthing accidents, congenital defects and infectious processes.

Wild mice are communal nesters and females often share duties in the care for the young^49^. This characteristic is maintained in laboratory mice and is often exploited by housing mice in breeding trios, which allows for increases in the number of pups per cage^47^ and individual pup growth^15, 16^. However, this social conformation is more prone to litter overlap^3^, the existence of a previous litter inside the cage, which is associated with higher neonatal mortality^3^.

There is a widespread opinion amongst laboratory mouse care staff that dams often kill their pups (infanticide) due to observations of entire litters disappearing or to finding partially eaten pups (cannibalism). Cannibalism is agreed to be a common occurrence in rodents, but actual observations of infanticide are rare^48^.

The occurrence of congenital malformations incompatible with life could also play a role in neonatal mortality and these are typically related to the strain. The most common congenital malformations in C57BL/6J mice are hydrocephalus and microphtalmia^4^, not directly related to neonatal survival.

C57BL/6J mice are the most used inbred strain in biomedical research, as a wild-type and also as background strain to produce genetically modified lines. Of the over 2.7 million genetically modified mice used in research in the EU in 2017^10^, the majority are expected to be on a C7BL6/J background. Identifying the major causes of perinatal mortality in this strain is crucial to develop preventive strategies for mouse breeding, but also in the context of phenotyping genetically modified mice, so that pathologies related to the background strain can be distinguished from those related to the genetic modification.

Neonatal mice are often not subject to a post-mortem analysis perhaps due to a lack of proper guidelines. To our knowledge, this paper provides the first systematic report of post-mortem findings in neonatal (0 to 4 days old) wild-type inbred mice found dead in laboratory animal facilities during routine inspection.

## Materials and Methods

### Ethical statement

Animals examined in this study originated from breeding colonies of healthy wild-type mice in two licensed laboratory animal breeding facilities. No animals were generated solely for the purpose of this study.

All breeding was done according to the UK and Portuguese legislation (Home Office Regulations (Animal Scientific Procedures Act 1986) and Decreto-Lei 113/2013) and Directive 2010/63/EU of the European Parliament following the 3R’s principle of Replacement, Reduction, and Refinement.

### Animals and housing

Breeding mice C57BL/6J were housed in two different animal facilities: F1 (i3S Animal Facility– Institute for Research and Innovation in Health, Porto, Portugal), and, F2 (Biological Support Unit – Babraham Institute, Cambridge, UK). Genetic integrity was maintained by breeding stock replacement every ten generations either by obtaining new stock directly from the supplier (The Jackson Laboratory) or rederivation from frozen embryos.

Each facility bred mice for research according to EU and national law governing animal research and had the necessary approvals. All pups collected were from litters from the regular breeding programs of the respective facility and no extra breeding occurred for the purposes of this paper.

F1 - Breeding pairs and trios were maintained under SPF conditions in conventional static cages (Pairs: 1264C Type II ©, Tecniplast, Italy, 268 × 215 × 141 mm; Trios: 1290D Type III ©, Tecniplast, Italy, 425 × 276 × 153 mm) with standard feed (Tecklad Global Diet Rodent 2014S, Italy) and water *ad libitum* obtained by reverse osmosis, filtered and UV irradiated. Each breeding cage was provided with a paper towel, a cardboard shelter (Des. Res.™, LBS Biotechnology, UK) and corncob bedding (Scobis duo, Mucedola, Italy). The room temperature was maintained between 20-24°C and relative humidity between 45-55% with a 12:12 light/dark photoperiod with no twilight.

F2 – Breeding pairs and trios were housed in individually ventilated cages (GM500 Filter©, Tecniplast, Italy; L × W × H, 391 × 199 × 160 mm), mounted on a ventilated holding unit (DGM Sealsafe Plus Rack©, Tecniplast, Italy). Cleaned cages were automatically dispensed with an average of 48 g of soft wood flakes bedding (EcoPure Chips 6 Premium©, Datesand, UK) and 7.5 g of white paper rolls (Enrich-n’Nest©, Datesand, UK) as nesting material. Cages had either a mouse loft (Tecniplast, Italy) or a red polycarbonate tunnel (International Product Supplies Ltd, United Kingdom; L × Ø, 98.55 × 50.80 mm) suspended from the grid lid, to provide in each case enrichment and a refuge in the event of a cage flood. Room temperature was maintained at 20–24 °C, relative humidity at 45–65% and a 12:12 h light regime with lights gradually switched on from 07:00. Standard food pellets (CRM (P) Vacuum Pack, Dietex International Ltd., UK) and filtered water obtained by reverse osmosis and ultraviolet irradiation were provided *ad libitum*.

### Pup collection and cage inspection

324 pups found dead were collected between May 2018 and September 2019 in Facility 1 (n=167) and between October 2019 and January 2020 in Facility 2 (n=157). Whenever a dead pup was found, it was collected, inspected, identified and put inside a plastic bag for later analysis.

In Facility 1, breeding cages were inspected daily for the occurrence of births and pup counting between the day the female was confirmed as pregnant through abdominal swelling until day four after birth. Cages were removed from the rack and opened inside a laminar flow cabinet. Pups were counted with minimal manipulation or nest disturbance.

In Facility 2 the procedure was the same with the exception of the adults being marked with a sterile surgical grade marker every two days (avoiding the day of parturition) as a part of a behavioural study on maternal behaviour to allow easy identification in video recordings, a procedure that lasted approximately 20 seconds for each animal.

In both facilities, cage inspection was performed by the same researcher at 10h ± 1h. Dead pups were also collected by technicians when found on daily inspection of breeding cages, using the same technique.

### Post-mortem inspection

Pups collected were refrigerated (n=171) or frozen (n=153) immediately until post-mortem analysis. These were defrosted at room temperature on the day the post-mortem analysis was performed. Fresh specimens were analysed within 4 hours of collection.

The necropsy protocol consisted of an external and internal inspection. The external inspection evaluated general condition, skin integrity and occurrence of acquired lesions such as bruises or wounds. Congenital abnormalities such as cleft palate, microphtalmia, alterations of head shape or limbs were recorded, as were the appearance of the umbilicus and presence of fragments of foetal membranes.

Internal inspection was performed under a stereomicroscope. This was initiated by performing skin incision along both abdominal and thoracic cavities; access to the abdominal cavity was gained by performing a circular incision along the abdominal wall. Attention was given to the presence and appearance of all internal organs and stomach contents were evaluated as well. To perform the inspection of the thoracic cavity, a portion of the thoracic wall was removed. Lungs morphology was evaluated and a lung fragment was removed in order to perform a lung float test, using the liver as a control. The results of this test could be correlated with the content of the stomach. The appearance of interscapular BAT (coloration, dimensions and condition of blood vessels) was recorded.

### Lung morphology and lung float test

Lung morphology was evaluated for degree of inflatedness, colouration and aeration. A lung float test was performed by removing a fragment of a lung lobe and a same size fragment of liver (as control for decomposition^32, 38^) and putting both inside an Eppendorf tube filled with water. Whether the fragments sank or floated was recorded.

### Evaluation of stomach contents

The level of distension of the stomach was evaluated and a small incision was made along the greater curvature. Stomach contents or lack thereof were recorded. Air was recorded whenever air bubbles where visible inside the stomach with or without other contents and some distension could be observed.

### Determination of stillbirths

Results of the lung float test were correlated with macroscopical analysis of the lungs, stomach contents and BAT.

A pup was considered a stillbirth when all the 8 criteria presented in Table 1 were fulfilled simultaneously. Any pup that did not meet all criteria were considered livebirth or, in the absence of any organ, not conclusive.

**Table 1.** Criteria for stillbirth evaluation.

| Table 1 - Criteria for stillbirth evaluation |
| --- |
| <u>Lung morphology</u><br>• No signs of aeration<br>• No signs of inflatedness<br>• Pale pink colouration |
| <u>Lung float test</u><br>• Sinking lung fragment<br>• Sinking liver fragment |
| <u>Stomach contents</u><br>• Presence of amniotic fluid inside stomach<br>• No air present |
| <u>BAT colouration</u><br>• Brown-yellow |

### Brown adipose tissue

BAT in the interscapular area was characterized in terms of its presence/absence, colouration (brown-pink or brown-yellow), presence of visible blood vessels and dimensions.

## Results

### Population characteristics

324 pups originating from a total of 154 litters of a population composed of breeding trios and pairs parities between 1 and 11 (Table 2).

**Table 2.** Population characteristics.

| **Table 2 – Population characteristics** |  |  |
| --- | --- | --- |
| **Breeding configuration** | **n** | **%** |
| Trios (2♀+1♂) | 114 | 74 |
| Pairs (1♀+1♂) | 40 | 26 |
| **Mortality** | **n** | **%** |
| Litters from which all pups died | 60 | 39 |
| Litters from which at least one pup survived | 94 | 61 |
| **Parity distribution** | **n** | **%** |
| Parity 1 | 11 | 7.1 |
| Parity 2 | 29 | 18.8 |
| Parity 3 | 37 | 24.0 |
| Parity 4 | 21 | 13.6 |
| Parity 5 | 23 | 14.9 |
| Parity 6 | 12 | 7.8 |
| Parity 7 | 10 | 6.5 |
| Parities 8-11 | 11 | 7.1 |
| **Parental age** | **Mean** | **SD** |
| Mother Age (days) | 165.9 | 56.8 |
| Father Age (days) | 175.9 | 55.8 |
| **Parental age range** | **Youngest** | **Oldest** |
| Mother Age Range (days) | 68 | 328 |
| Father Age Range (days) | 77 | 317 |

*n* - number of occurrences. % - Percentage of occurrences. Mean - arithmetic mean. *SD* – Standard deviation. ♀ - Female. ♂ - Male

### Lung evaluation

Lung float tests were performed in 321 pups. 65% of the lungs tested floated and 35% sank. All control liver fragments sank.

Lung colouration was generally pale pink for sinking lungs, and pink or dark pink for floating lungs, the latter possibly related to hypostatic blood accumulation. Since dead pups were collected whenever found, decomposition could be expected in some specimens. These commonly partially floated but all had signs of aeration and inflatedness with observable air bubbles in the pulmonary parenchyma.

Two pups presented severely congested lungs with a grey colouration and the lung fragment sank, although some level of lung inflation or aeration was observed. One of these pups also showed blood clots inside the thoracic cavity.

Three pups were not evaluated due to missing lungs (cannibalism).

### Stomach contents evaluation

Stomach contents evaluation of all pups are summarised on table 3.

**Table 3.** Stomach contents.

*n* - number of occurrences. % - Percentage of occurrences. Mean - arithmetic mean. *SD* – Standard deviation. ♀ - Female. ♂ - Male
| Table 3 – Stomach contents |  |
| --- | --- |
| Contents | n(%) |
| Air | 138<br>(42.6%) |
| Amniotic fluid | 99 (30.6%) |
| Empty | 69 (21.3%) |
| Decomposed | 9 (2.8%) |
| Milk | 8 (2.5%) |
| Blood | 1 (0.3%) |
| Meconium | 1 (0.3%) |
| Fur | 1 (0.3%) |
| Missing | 10 (3.1%) |

*n - Number of pups. % - Percentage of pups*

Only 8 (2.5%) pups had milk in their stomachs and only two pups had a stomach filled with milk. Only one of these, a pup from a cage without an older litter, presented traumatic lesions (neck echimosis and bruises in the hindlimbs).

### Stillbirths and Livebirths

Overall percentages of stillbirths and livebirths are illustrated in Figure 1. Seventy (21.6%) pups were considered stillbirths, according to the criteria in Table 1. Of these, 17 (24.3%) also presented amniotic fluid inside the thoracic cavity, 13 (18.6%) presented traumatic lesions of which one was partially eaten. Internal haemorrhage was observed in 2 pups and possibly was related to trauma during birth. 2 pups presented intact umbilical cords (Figure 2) and 6 pups presented abnormally small size (Figure 4).

**Fig. 1.**
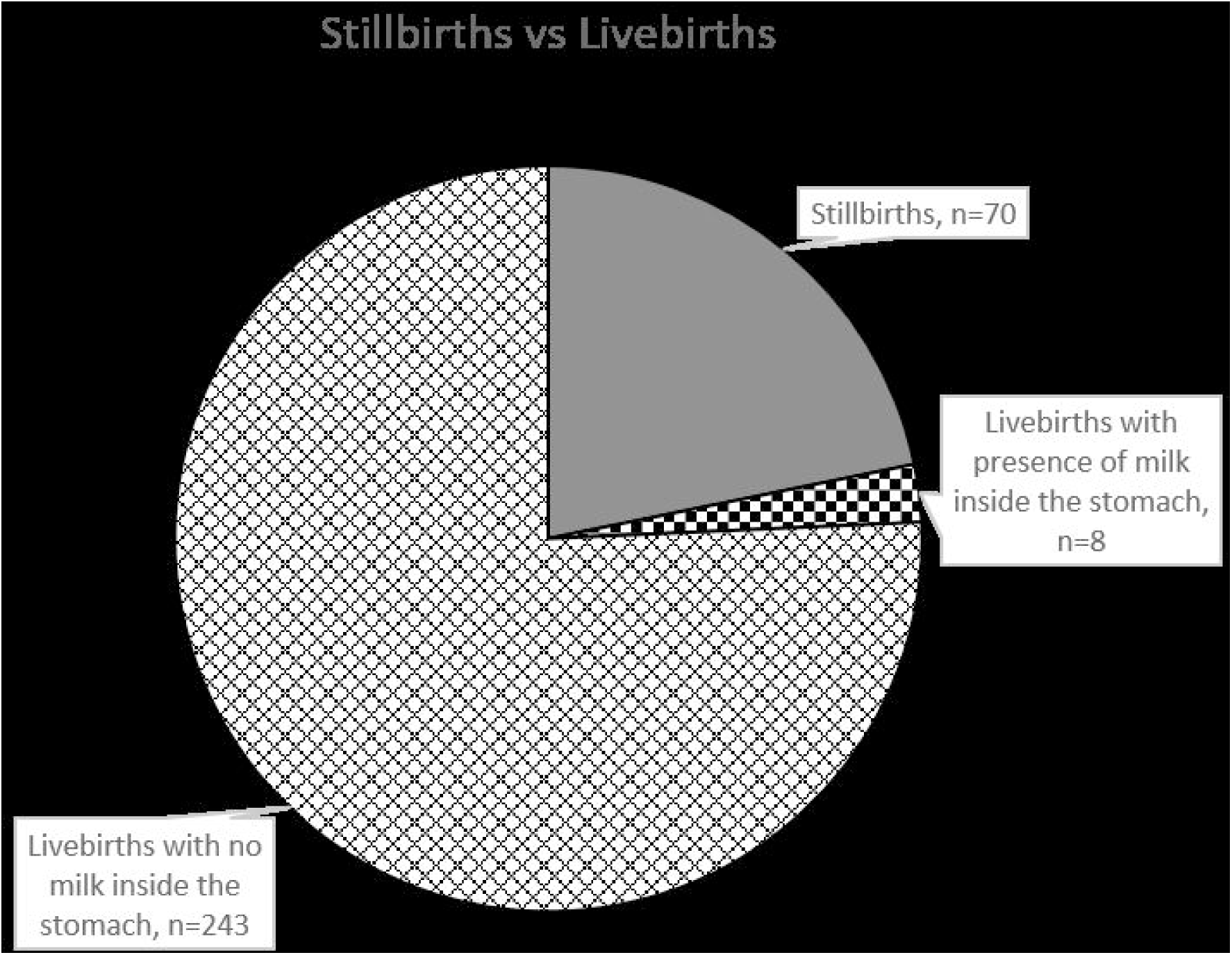
Numbers of stillbirths and livebirths with and without presence of milk inside the stomach.

**Fig. 2.**
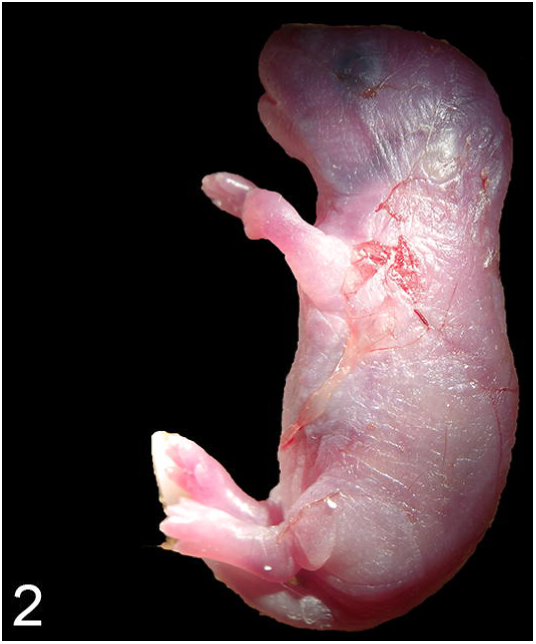
Remains of foetal membranes and umbilical cord adhered to the skin.

251 (77.5 %) of the pups inspected were considered livebirths, with all lungs showing some degree of aeration or inflation. These pups were found 0,8 ± 0,83 days after birth. Stomach contents were: milk (n=8; 3.2%), amniotic fluid (n=29; 11.6%), air (n=133; 53.0%), empty (n=69; 27.5%) and decomposed or stomachs missing (n=16; 6.4%). Traumatic lesions were observed in 63 (25.1%) of livebirths and 18 (7.2%) were partially eaten. Internal bleeding was diagnosed in 14 pups by observation of blood inside the abdominal cavity.

3 (0.9%) pups could not be evaluated due to cannibalism as some of the internal organs (including the lungs) were missing.

### Brown adipose tissue

BAT was present in all of the pups inspected. The tissue colouration, blood vessel and dimensions varied according to the age of the pups. On day 0 pups usually presented brown-yellow BAT with visible blood vessels, by day 1, brown-pink BAT was observed with variably reduced appearance of blood vessels.

### Pathological findings and traumatic lesions

All gross pathological findings are summarized on Table 4. Overall, 79 (24.4%) pups presented some kind of traumatic lesion, including bite wounds and bruises (Figure 5-7). Bite wounds include cannibalism events where parts of a pup were missing and small incised or puncture wounds probably caused by biting.

**Fig. 5.**
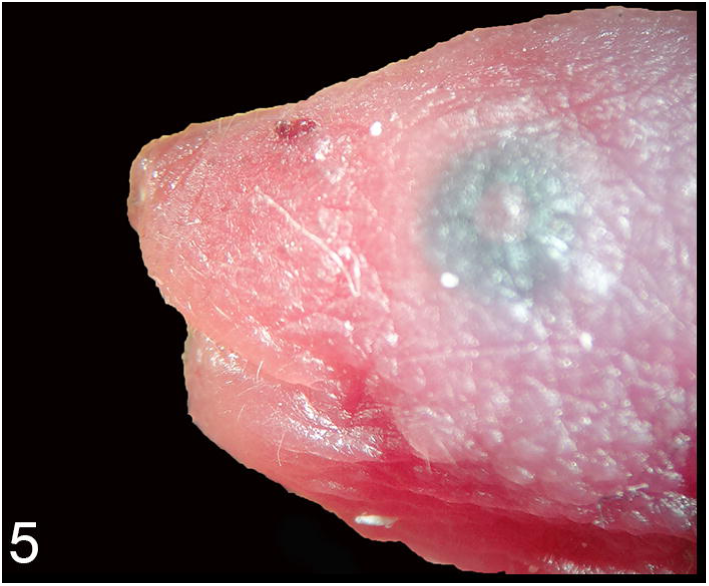
Small wound left dorsal area of muzzle.

**Fig. 6.**
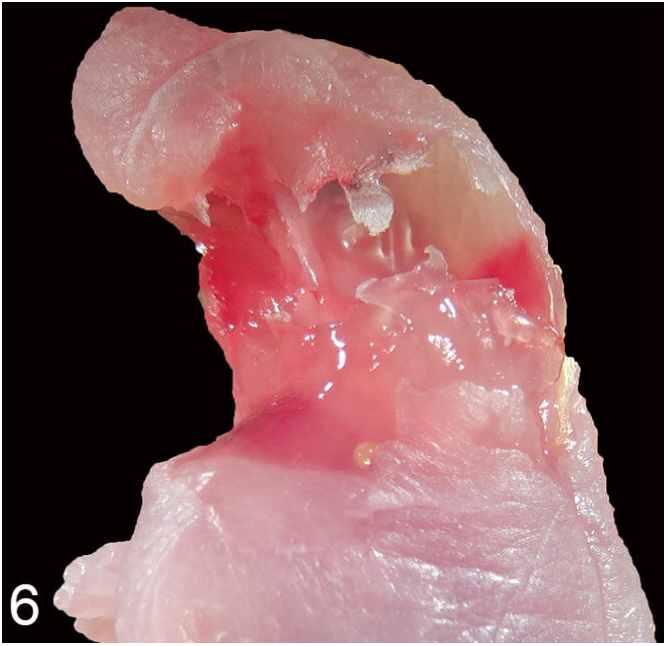
Cannibalism. Left portion of cranium and neck missing.

**Fig. 7.**
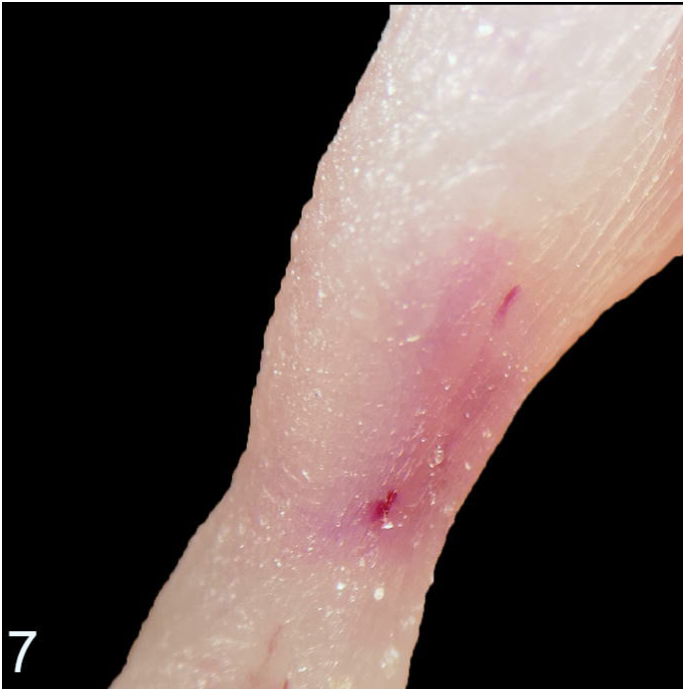
Two small puncture wounds medial area of right hindlimb.

**Table 4.** Pathological findings and traumatic lesions distribution.

*n - Number of pups. % - Percentage of pups*
| **Table 4 – Pathological findings and traumatic lesions distribution** |  |  |
| --- | --- | --- |
| **Pathological findings** |  |  |
| **Organ** | **Finding** | **n(%)** |
| Heart | Cardiomegaly | 1 (0.3%) |
| Liver | Hepatomegaly | 1 (0.3%) |
|  | Colour alterations | 60 (18.5%) |
| Thymus | Petechiae | 3 (0.9%) |
| Eye | Microphthalmia | 1 (0.3%) |
| Spleen | Colour alterations | 5 (1.5%) |
| Lungs | Congestion | 2 (0.6%) |
| Skin | Congenital defects (localized) | 2 (0.6%) |
| Systemic | Subcutaneous edema (Anasarca) | 2 (0.6%) |
|  | Hydrothorax | 2 (0.6%) |
|  | Ascites | 2 (0.6%) |
| **Traumatic Lesions** |  |  |
| **Type of lesion** | **Localization** | **n(%)** |
| Bruises | Head | 23 (7.1%) |
|  | Limbs | 11 (3.4%) |
|  | Abdomen | 4 (1.2%) |
|  | Thorax | 2 (0.6%) |
|  | Dorsal area | 3 (0.9%) |
| Bite wounds | Head | 21 (6.5%) |
|  | Limbs | 10 (3.1%) |
|  | Abdomen | 11 (3.4%) |
|  | Thorax | 1 (0.3%) |

205 (63.3%) pups were collected from cages where there was an older litter present, from these 23.4% (48) presented traumatic lesions. While from the group of pups that were collected from cages with no previous litter (n=119; 36.7%), 26.1% (31) of these showed traumatic lesions. Excluding bite wounds, the incidence of traumatic lesions such as bruises in the overlap group was 20% (41) and 16% (19) in the no overlap group.

Regarding stomach contents, in the overlap group, 5 (2.4%) pups presented milk inside stomach and 140 (68.3%) presented empty stomachs or presence of air inside the stomach. In the no overlap group, 3 (2.5%) pups presented milk inside the stomach and 52 (43.7%) presented empty stomachs or presence of air inside the stomach.

Lesions involving structures of the head such as the lips, jaw or tongue were present in 9 (3.6%) pups considered live-born. Internal haemorrhage was seen in a total of 13 (4.0%) pups with significant amounts of blood within the abdominal cavity; of these 11 had evidence of being live-born.

Hepatomegaly and cardiomegaly were seen in one case. Two pups presented anasarca (Figure 3), hydrothorax and ascites.

**Fig. 3.**
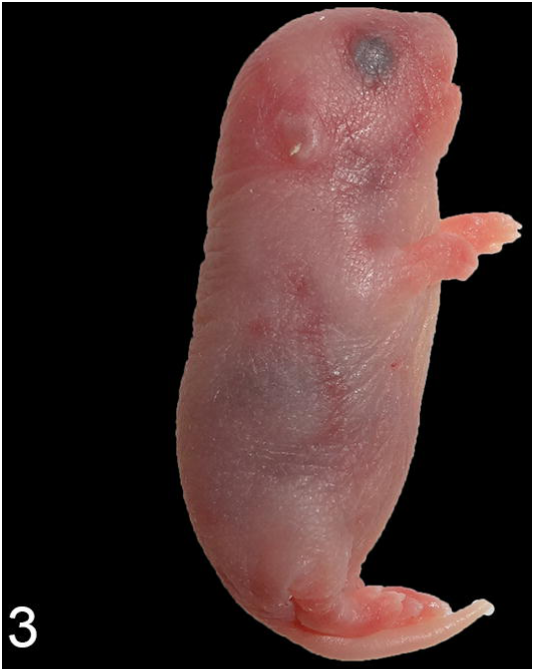
Anasarca.

**Fig. 4.**
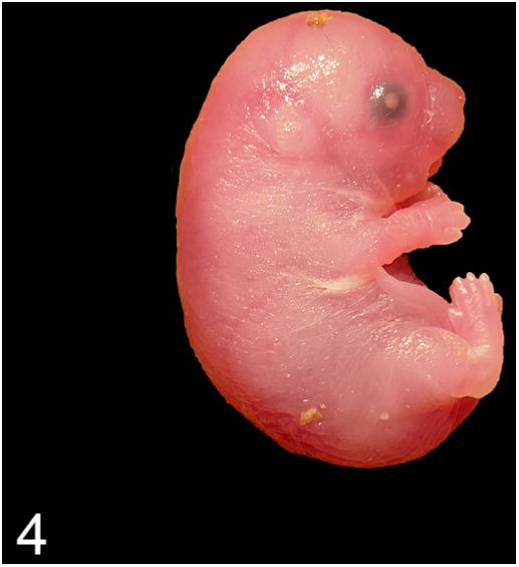
Stillbirth with kyphosis and abnormally small size.

### Congenital malformations

No pups inspected showed palate malformations. Microphtalmia was observed in one pup. Hydrops fetalis was observed in 2 pups (0,6%) which showed anasarca, hydrothorax and ascites (Figure 3). Skin defects localized in the head and the umbilical region were observed in two pups (0,6%) (Figure 8).

**Fig. 8.**
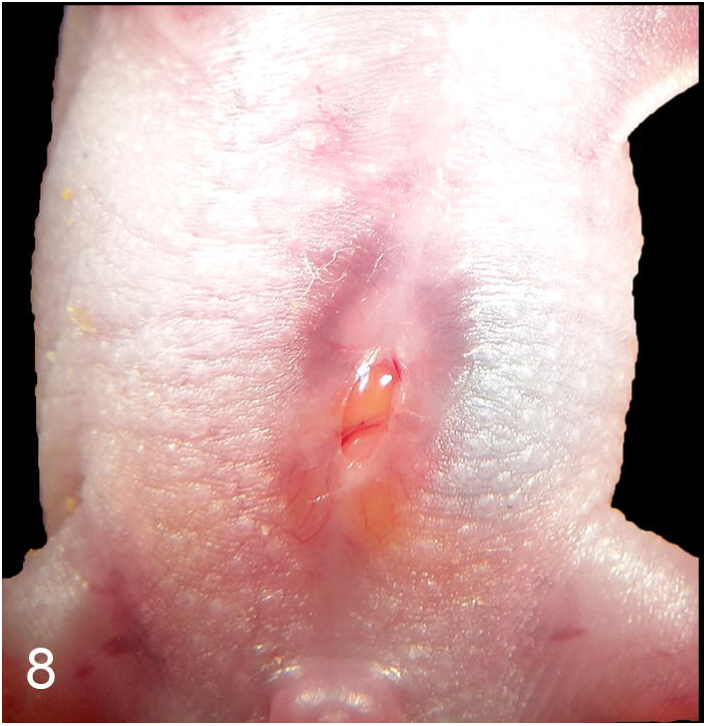
Skin defect at the umbilical region with exposed abdominal viscera.

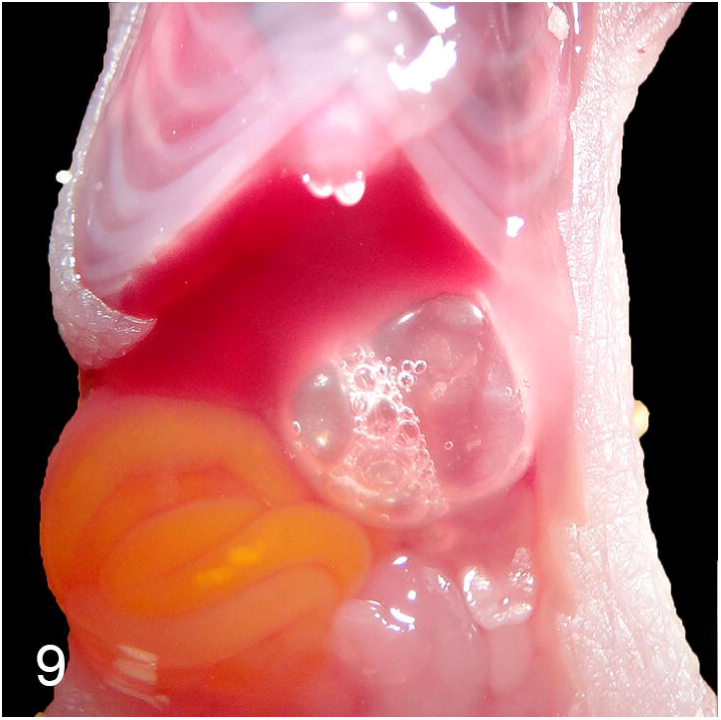

**Fig. 10.**
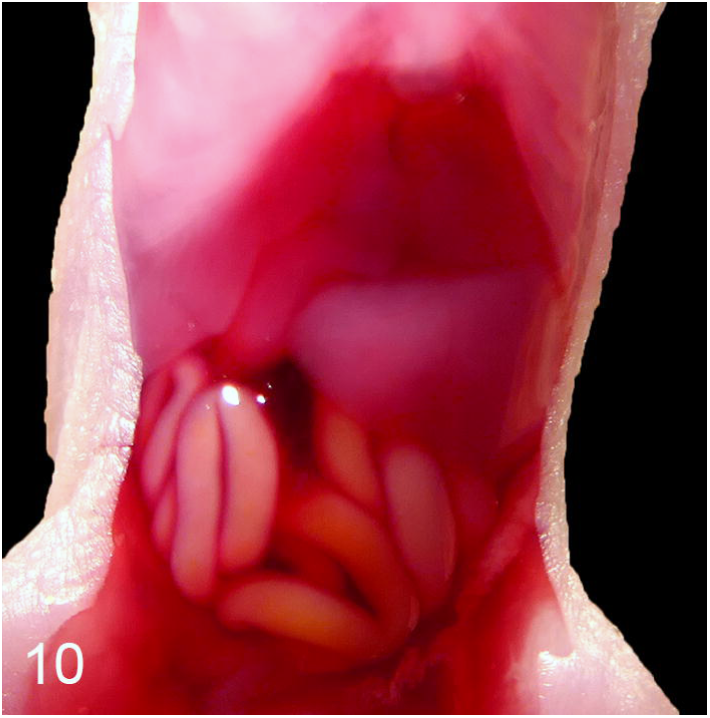
Hemoabdomen.

## Discussion

In this study, 21.6% of the pups found dead were considered stillbirths and 77.5% livebirths. Among the livebirths, only 3.2% presented with milk inside the stomach, indicating successful suckling. Congenital malformations were very rare, whereas traumatic lesions were found in 24.4% of all the pups.

One hypothesis for neonatal death of live born pups would be that impairments to maternal behaviour could compromise suckling. Neonatal pups, as all mammals, are dependent on the ingestion of colostrum and milk for survival, Without being fed, neonates are only able to survive for 24 hours^30^. The first suckling event and a successful nipple attachment is therefore critical. For this, mutual dam and pup recognition is essential. This process seems to be mediated by odour cues that are present in the nipples of the dam and are recognized by the pups^1^. Since pups are relatively immobile at birth maternal attention is required for a successful nursing event. Ultrasonic vocalizations (USVs) play a major role in the communication between the dam and pups and also in the formation of the mother-infant bonding along with olfactory and visual stimuli^36^ so it is likely that they will also play a major role in pup survival.

Pups are highly vocal in the first two weeks after birth, producing audible calls in situations such as rough handling^13^ and, most notably, USVs that are emitted by pups when in distress (isolation and cold stress) eliciting a maternal response from the dam, such as pup retrieval^7^. These are characterized by having frequencies between 30-90 kHz and being emitted in an increasing rate during the first 6-7 days after birth and gradually decreasing until complete disappearance at two weeks after birth^8^. USVs emission is needed to ensure successful communication and, thus, a maternal response. Further evidence of their role in pup survival is given due to the finding that an increase in serum prolactin levels occurs as a response to USVs in lactating rats^14^.

C57BL/6 mice are known to suffer from age-related hearing loss that starts to develop as soon as 10 weeks of age and initially affects hearing of higher frequencies^20^. Furthermore, sex differences have been reported with females presenting a more severe degree of hearing loss when compared to males of the same age^17^. This hearing impairment can potentially have an impact on maternal care and compromise pup survival by decreasing maternal responsiveness and predisposing to death by hypothermia or starvation. Further investigations are needed on this topic.

More than half (53,4%) of livebirths presented air inside the stomach, either as the only content (50,1%) or associated with milk (3,3%). This finding could suggest that these pups tried to suckle but only air was ingested. Air inside stomach takes volume and could also impair the ingestion of milk and lead to malnutrition. The attempt to suckle and the ingestion of air suggests either a deficient lactation (maternal cause) or a failed attempt to nurse (neonatal cause).

For a successful attempt to suckle, pups need the necessary structures to be able to attach to the nipple and create negative pressure to ensure milk flow and ingestion. A failed attempt to suckle could be related to a deficient nipple attachment due to craniofacial deformities such as hard palate abnormalities^39^ or ankyloglossia^35^ that are generally associated with severe gastric distension. In this study, hard palate deformities and tongue immobility were not observed but further histologic analysis could be necessary to evaluate the presence of ankyloglossia. Furthermore, only two pups presented severe gastric distension.

A deficient lactation could also play a role. Milk production in mice can be influenced by several factors, including nutritional status^5^, food intake^22, 44^, temperature^12, 21, 26-28, 46^, litter size^24, 25^ and lactation number^24^. Pup stimulus after birth seems to play a major role since in previous studies differences in milk production were only found after the fourth day of lactation across different litter sizes^25^

In situations of litter overlap, an increase in temperature inside the nest can have a negative effect in milk production^12^. Furthermore, since female mice are known to nurse pups indiscriminately in communal nests^11^, it is possible that milk “theft” could occur due to competition from larger and more mobile older pups. However, our data show that the proportion of pups without milk in their stomach is the same for cages with overlapping litters as for cages with only one litter.

Infanticide is very rare in B6 mice^48^, suggesting that the traumatic lesions observed should not generally be regarded as the cause of death. Bruises in different locations of the body were observed at a low frequency (13%) varying in colour between red and blue, and sometimes associated with swelling. The timing of such lesions is difficult to determine but since their formation requires blood flow and some time for the lesion to appear, forensic pathologists agree that they should be normally regarded as indicators of an ante-mortem traumatic event, although it can happen post-mortem when greater force is applied^42^. Further histological and biochemical analysis could be helpful in timing these lesions^31^. However, the majority of the observed bruises were insufficient to be regarded as the cause of death, with the possible exception of those involving the lips, tongue or jaw that could have an impact on the ability to suckle. Of course, pups that have died and been entirely consumed are missing from this data set.

Internal haemorrhage was a rare observation in both live and stillbirths. In both cases, trauma would be required and blood loss would contribute to death. The occurrence in stillbirths is probably related to parturition issues^45^.

Congenital malformations such as cleft palate, hydrocephalus or other gross malformations in internal organs that could impair survival were not directly observed, although a skin defect around the umbilical area that could potentially expose internal organs and compromise survival was observed in one pup (Figure 8). Another pup presented signs of cardiac failure at 2 days of age (cardiomegaly and hepatomegaly possibly related to a cardiovascular malformation. Two pups presented anasarca, hydrothorax and ascites, all lesions compatible with fetal hydrops^37^. Microphtalmia was also observed in one pup.

Cannibalism is common in laboratory mice and infanticide has been attributed as a cause of neonatal death in laboratory animal house facilities^43, 48^ although our recent data suggests this is less common than previously reported^3^. In the present study, only one pup from 324 was clearly the victim of trauma ante-mortem sufficient to cause death by the presence of incised wounds, facial bruises with visible oedema and blood inside the stomach.

Identification of stillbirths has always been a challenge^40^ and the lung floatation test is the subject of debate^38^ as tissue decomposition can lead to confounding effects. Fortunately, in laboratory animal house facilities environmental conditions are highly standardized and the animals are inspected at least once every day. Therefore decomposition processes are limited and the rate relatively standardised. A lung float test and evaluation of stomach contents to identify stillbirths in breeding transgenic mice has been reported^9^. However, performance of the lung float test alone can lead to erroneous conclusions^34^. We therefore used the liver as a control for air produced by decomposition^32^. When combined with a careful evaluation of the lung morphology, stomach contents and brown adipose tissue, a reliable predictor of post-partum viability may be obtained. Further evidence could be obtained by observation of the umbilical cord (cicatrisation versus the presence of an intact umbilical cord and/or foetal membranes). Presence of amniotic fluid inside the stomach indicates either a stillbirth or a death immediately after birth because foetuses swallow amniotic fluid while inside the uterus^23^ but gastric emptying occurs as much as twice per hour in neonates^41, 50^.

In summary, the findings presented here show that the commonest cause of death in neonatal C57BL/6 laboratory mice is starvation since the majority of livebirths had no milk in their stomachs. Secondary reasons for a failure to suckle should be a key subject for future research. Stillbirth was the second most common reason for death, although the causes behind this could not be identified in most cases. Contrary to expectations, active infanticide by adults was extremely rare.

## Acknowledgements

The authors thank all staff at the i3S Animal Facility and the Babraham Institute Biological Support Unit, especially Ana Abreu, Ângela Ribeiro, Urszula Karpinska and Minnie Eve, and the i3S Histology and Electron Microscopy Platform, member of the national infrastructure PPBI - Portuguese Platform of Bioimaging (PPBI-POCI-01-0145-FEDER-022122).

## Supplemental Files

Original files of the post-mortem pictures are available on Supplemental File 1.

