## Supplementary figures and images for "Causes of death in newborn C57BL/6J mice"

### Supplemental File1

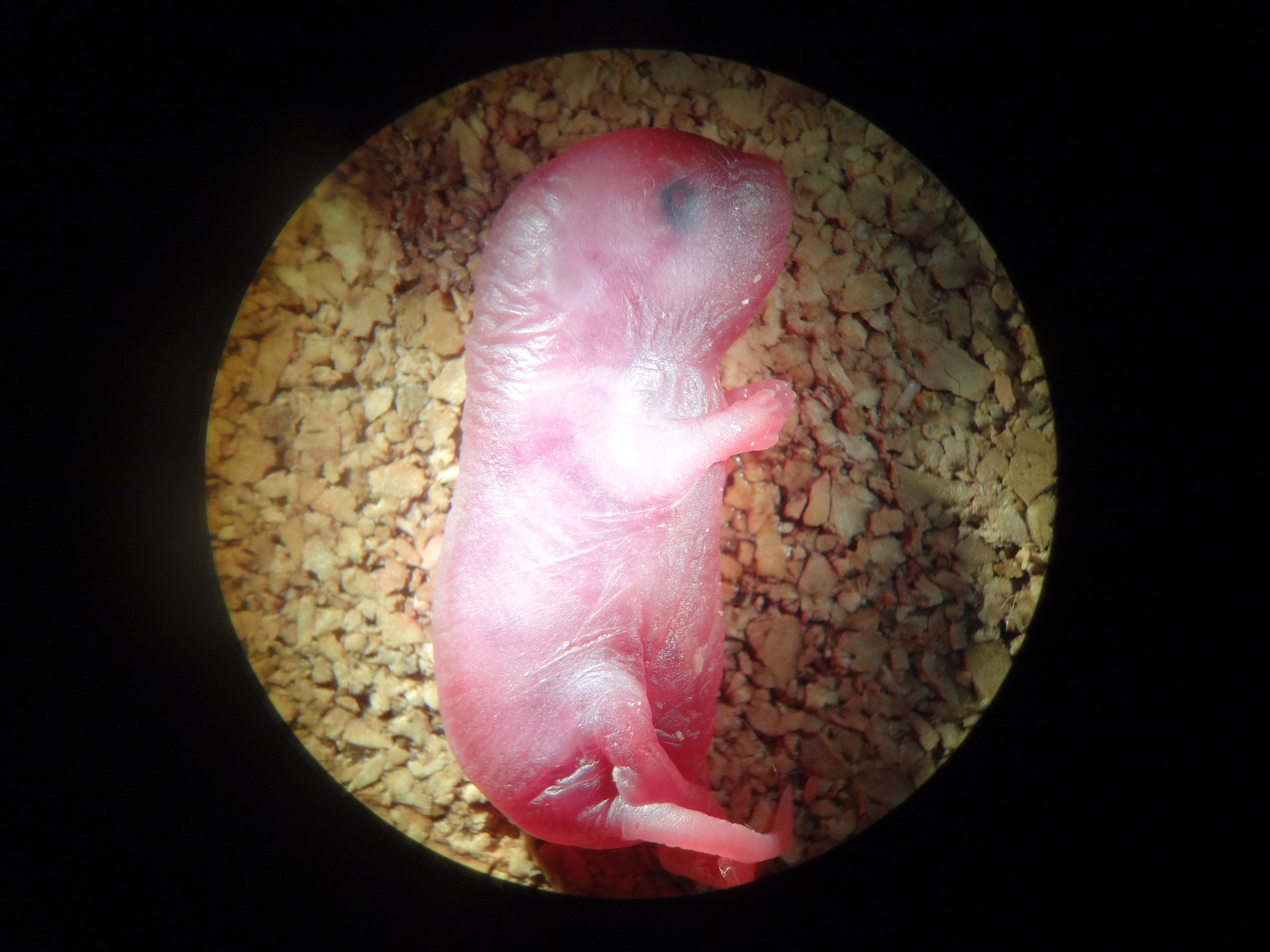

### Supplemental File1

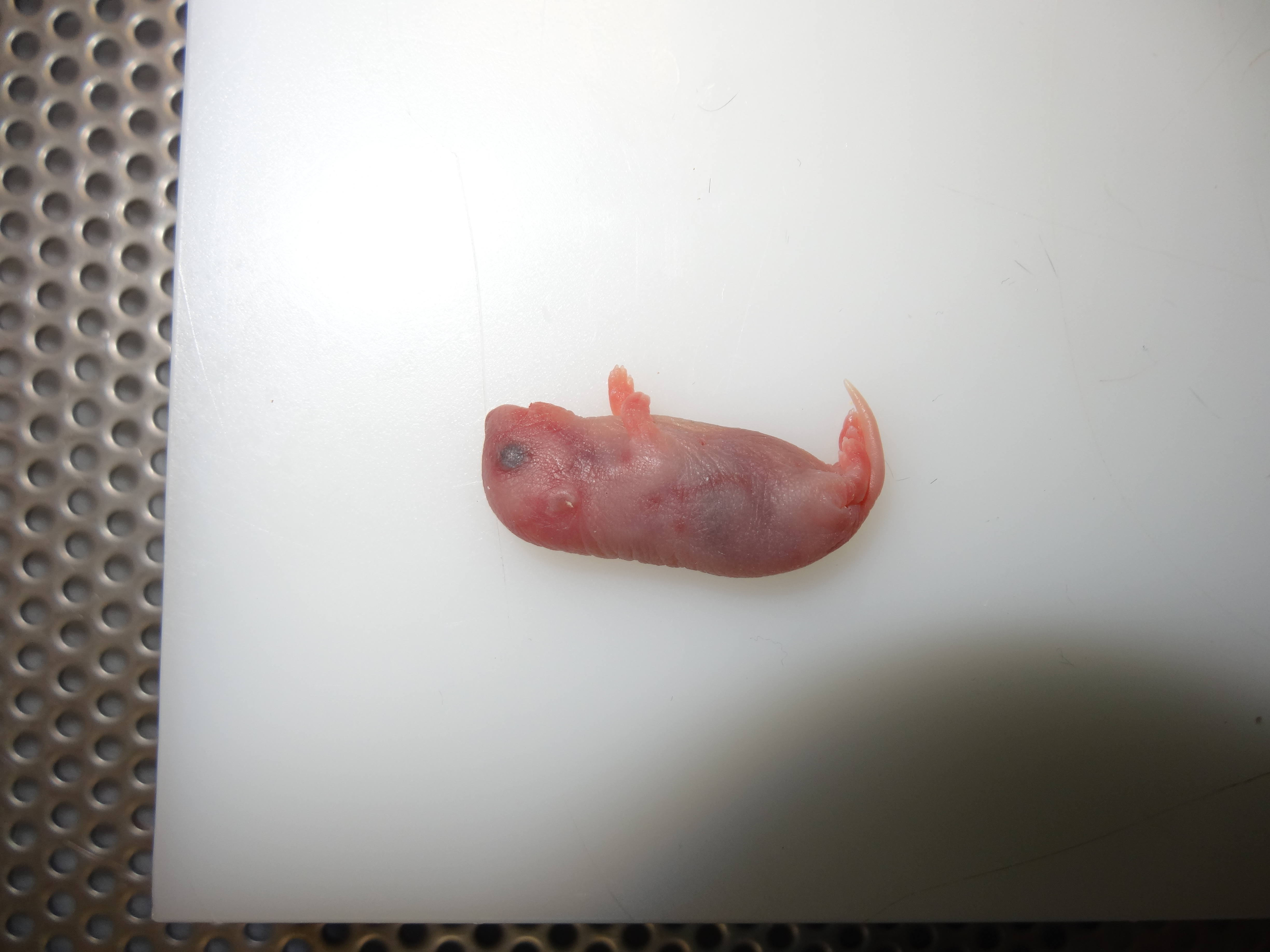

### Supplemental File1

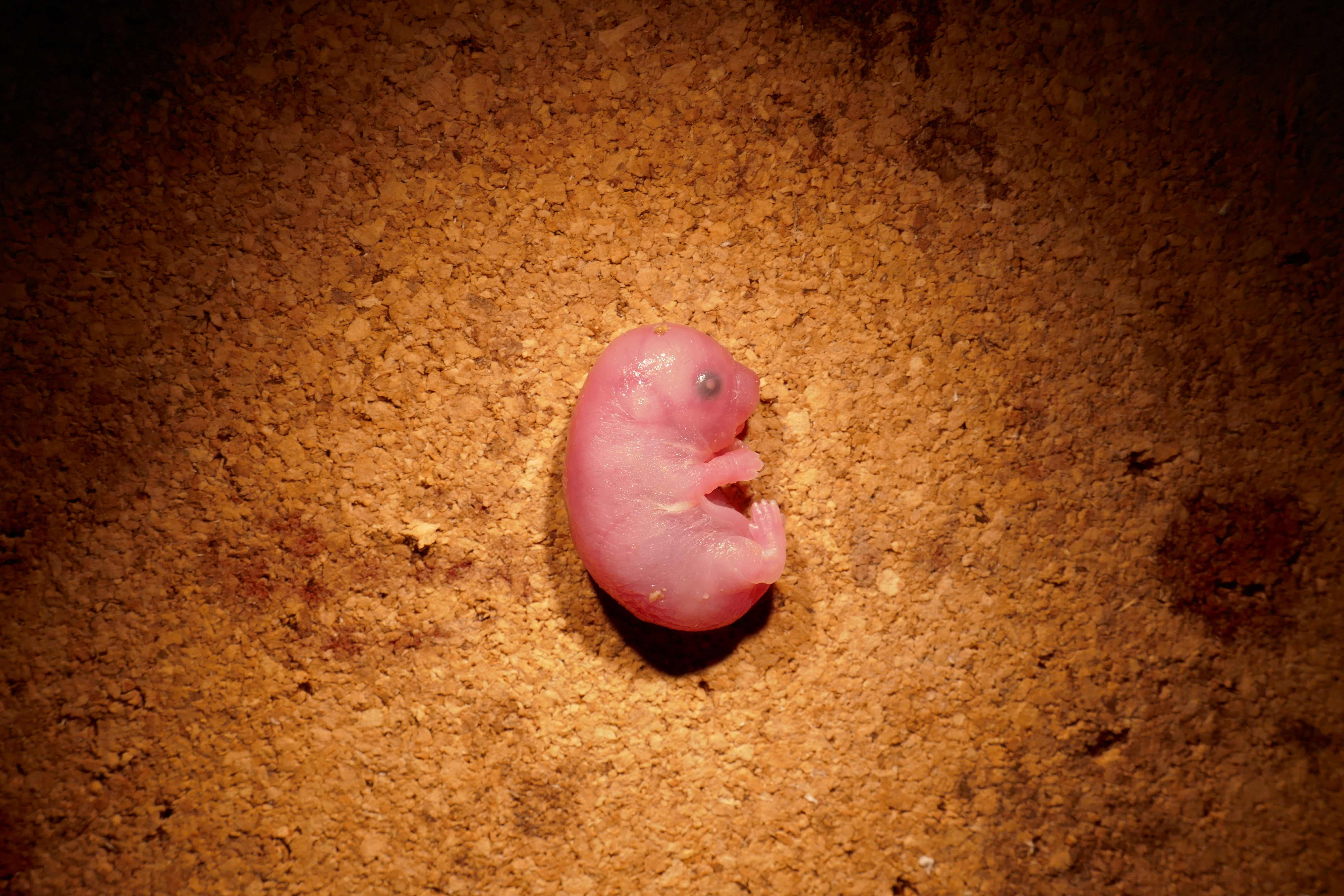

### Supplemental File1

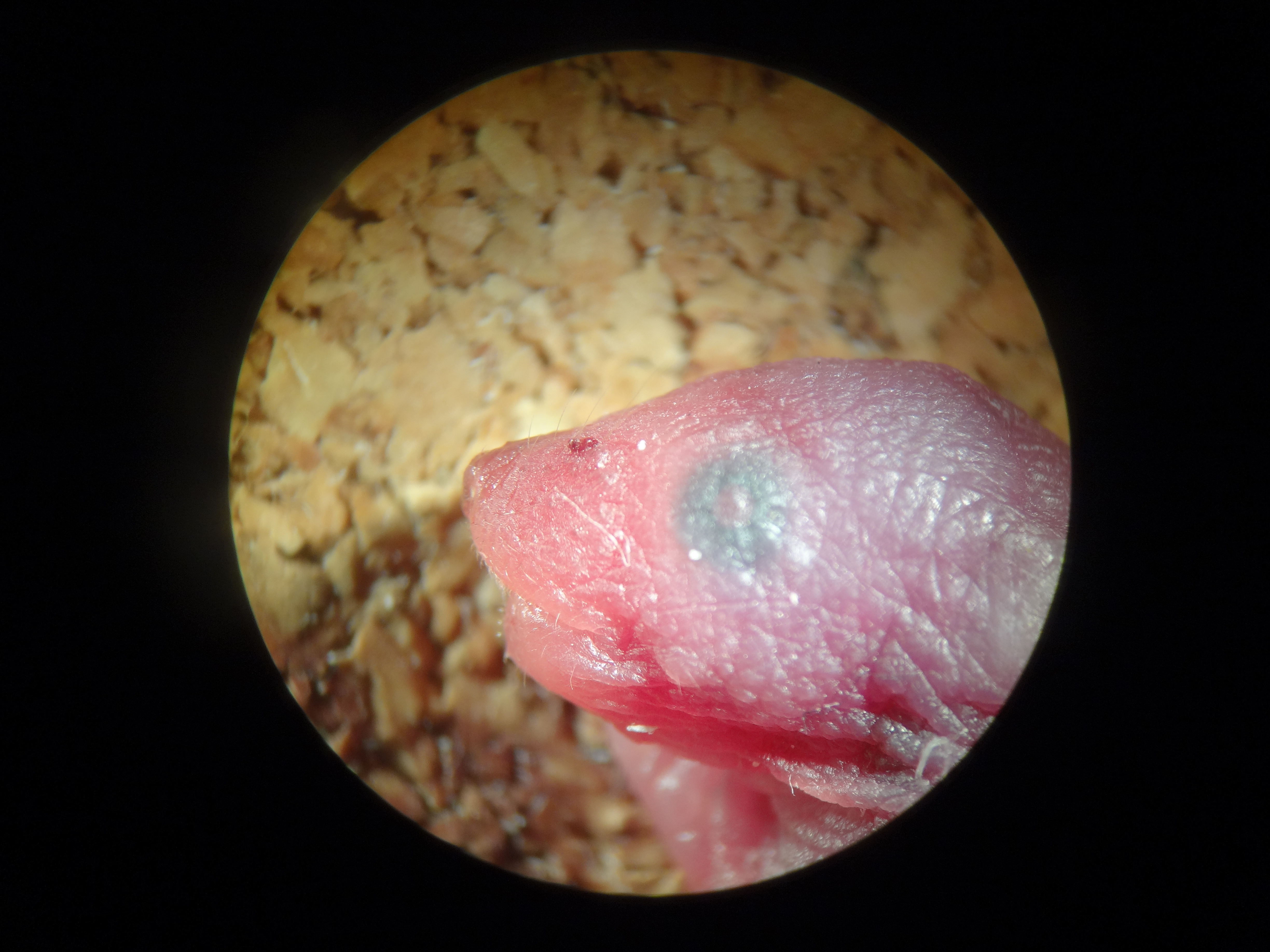

### Supplemental File1

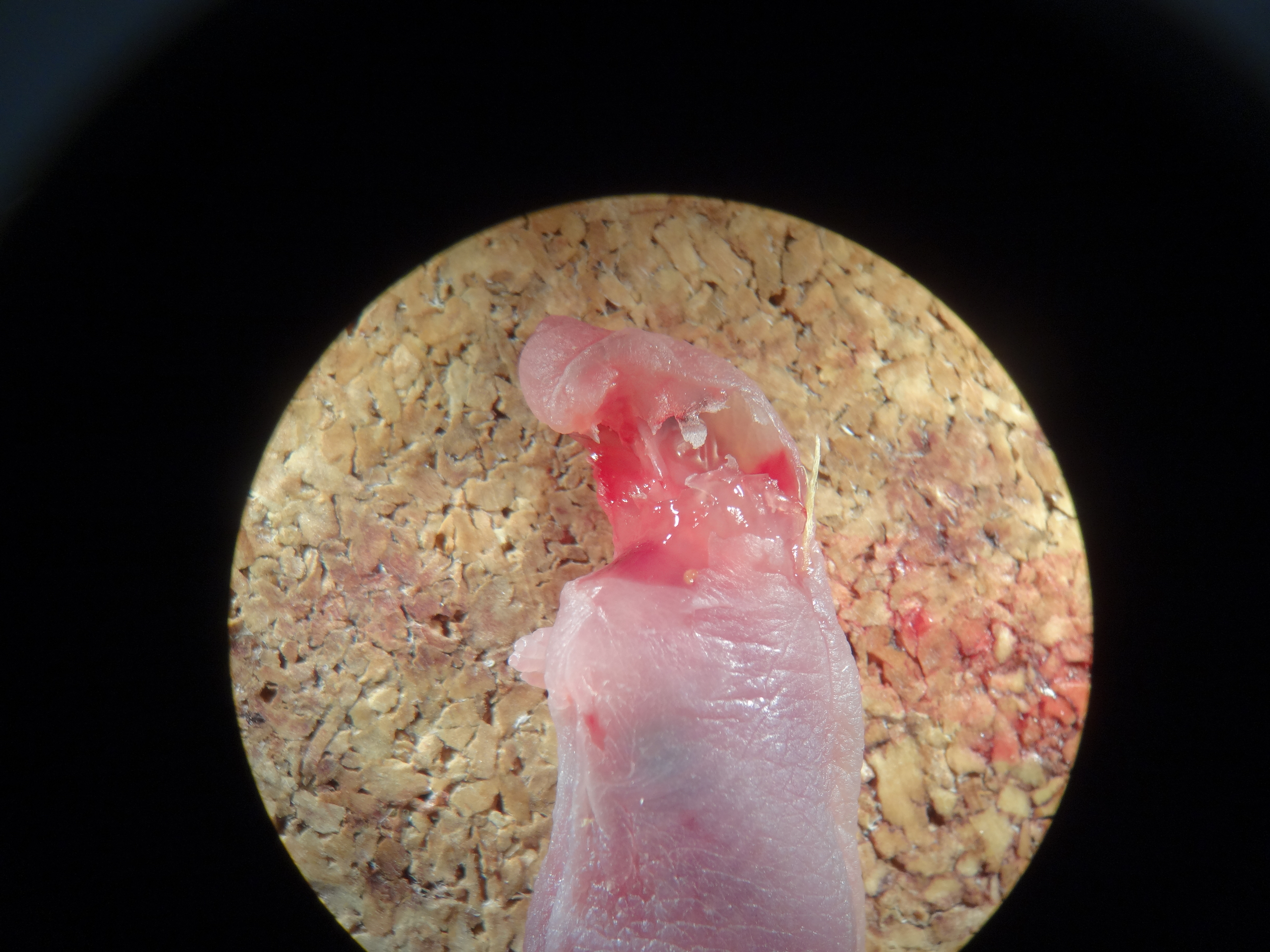

### Supplemental File1

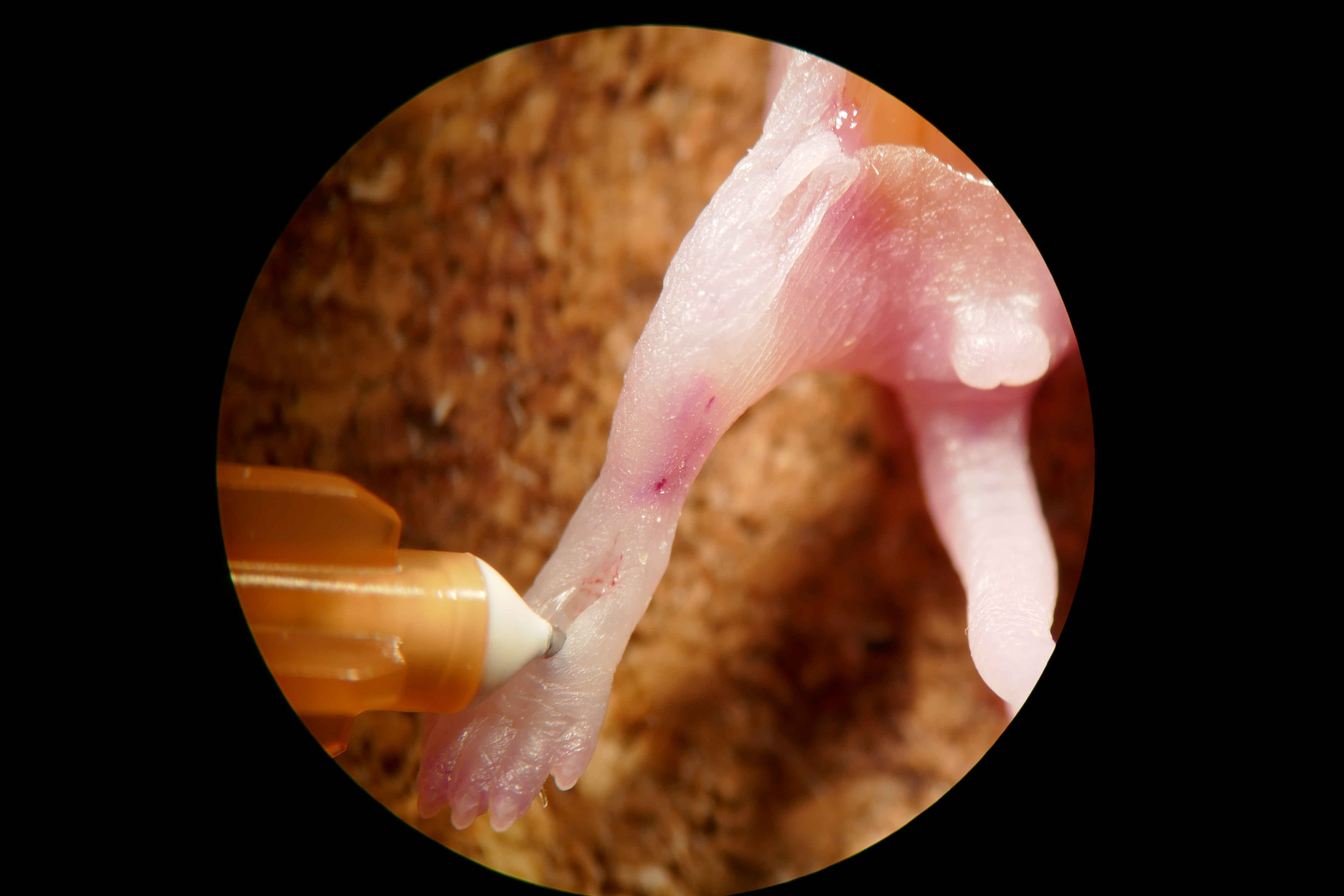

### Supplemental File1

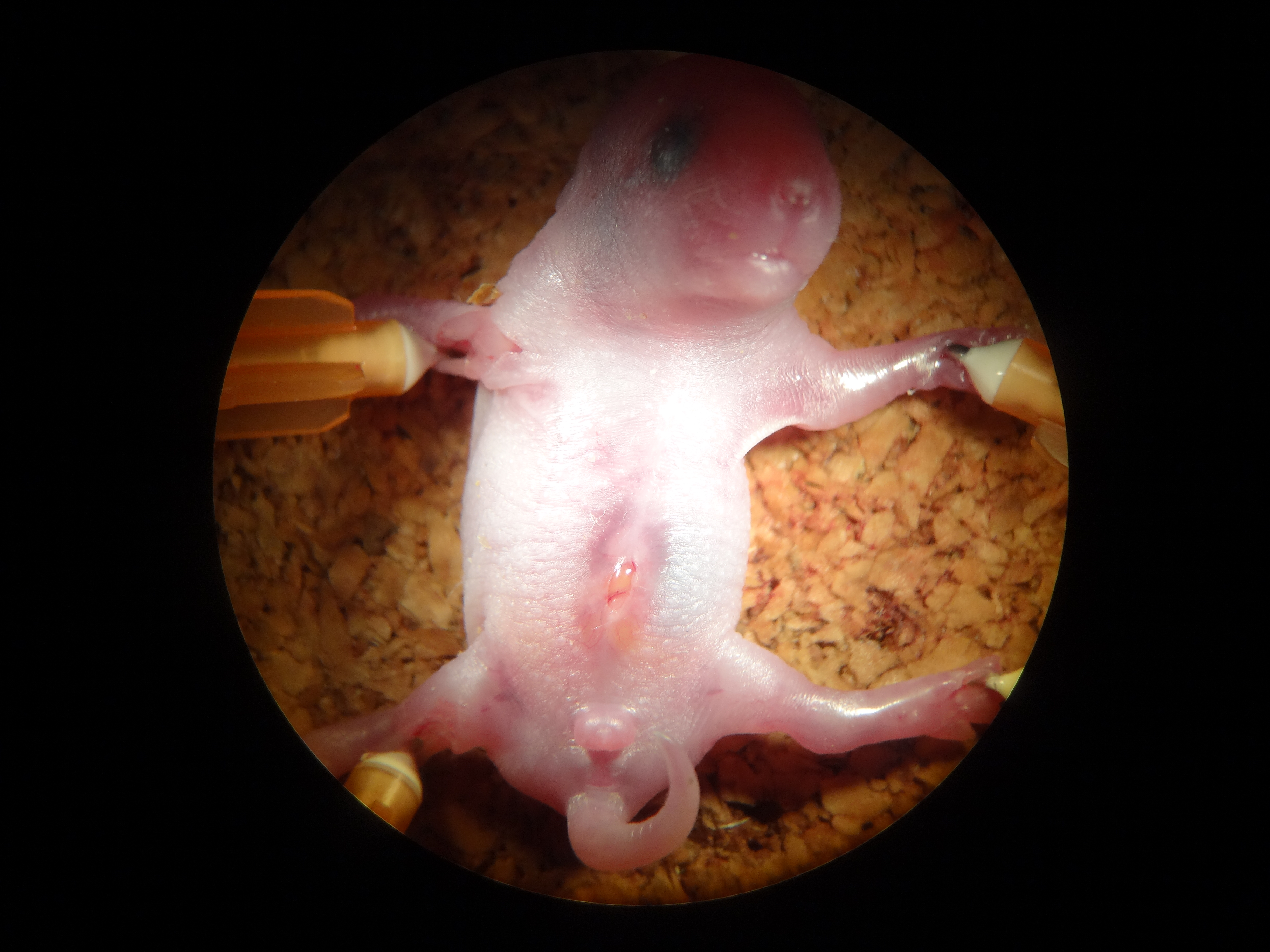

### Supplemental File1

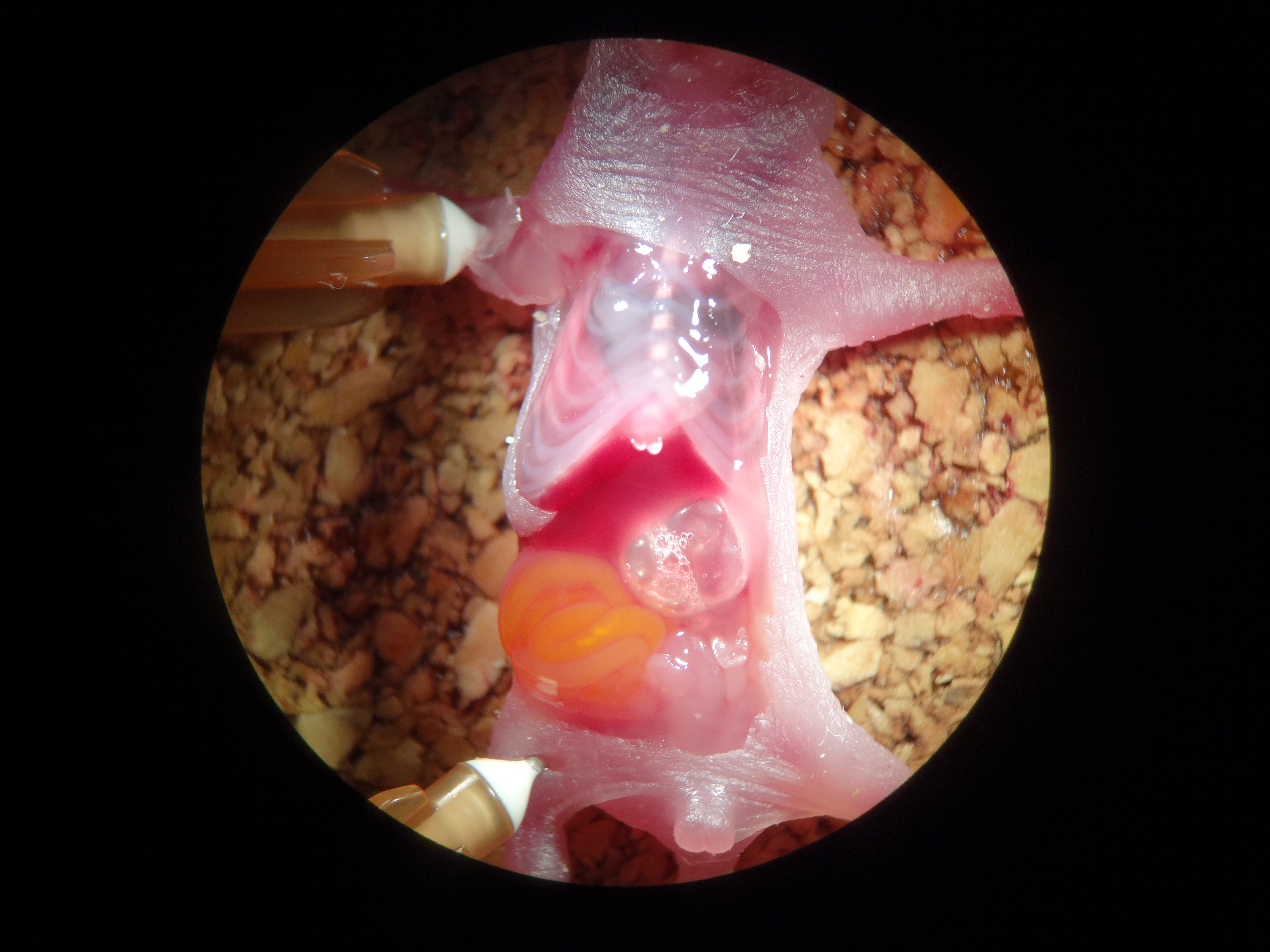

### Supplemental File1

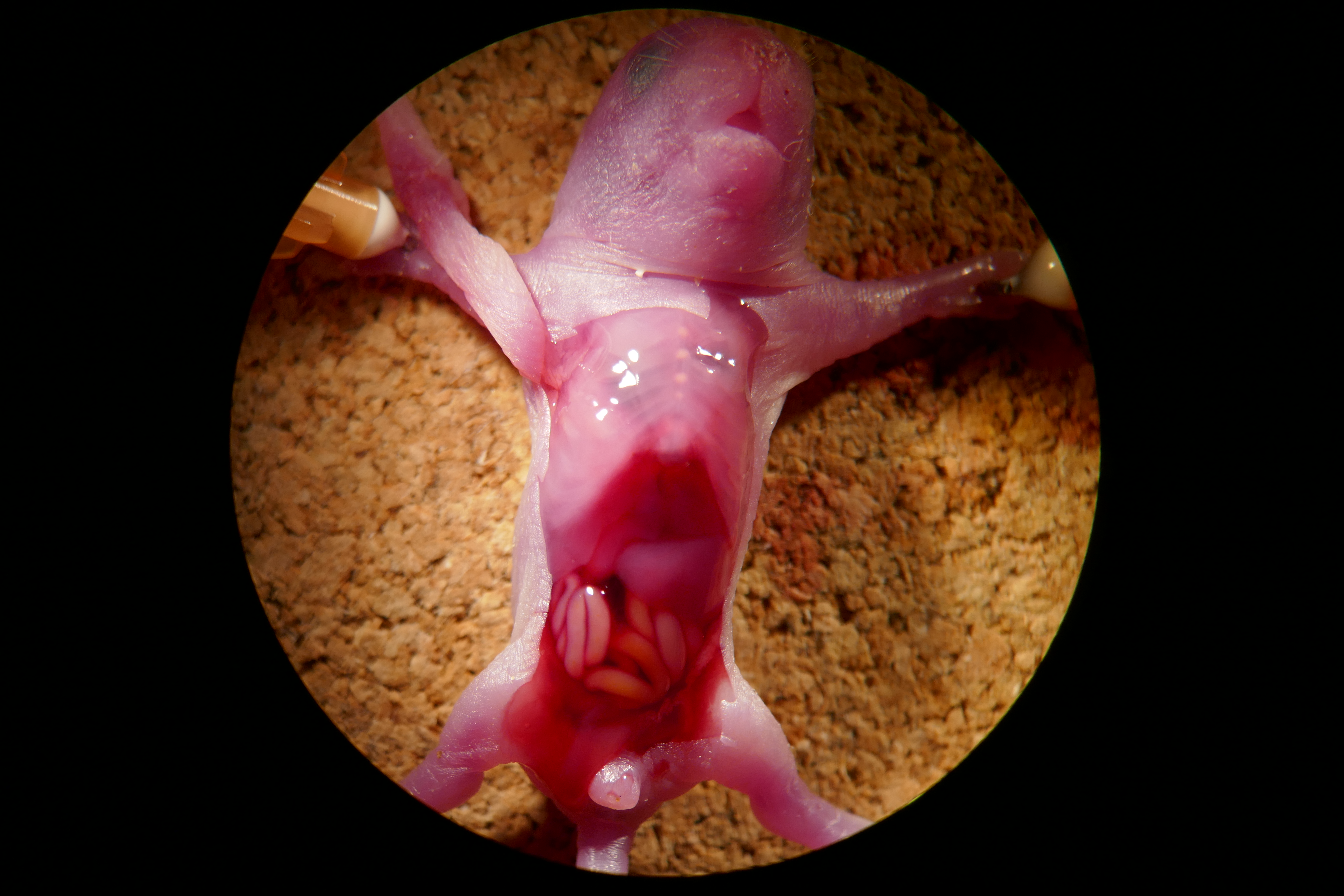
